# Avian cranial evolution is influenced by trade-offs in shape between hard and soft tissue traits

**DOI:** 10.1101/2025.03.21.644529

**Authors:** Andrew Knapp, Taylor West, Catherine M Early, Ryan N Felice

## Abstract

Changes in the structure and relative size of the brain are thought to be key transformations in the origins and continued evolution of birds, reflecting innovations and diversity of neurosensory and cognitive capabilities. However, these neuro-anatomical and functional changes do not occur in isolation, being accompanied by a host of other derived morphological characteristics associated with the evolution of flight. In the avian head alone, these include the evolution of a toothless beak, increase in relative eye size, and reduction and restructuring of jaw muscles. Several hypothesized developmental trade-offs have been proposed to explain the interrelationships among the hard and soft tissues of the head. How these developmental patterns translate into evolutionary trade-offs in other cranial traits is poorly understood, despite brain shape evolution being well documented in birds. Here, we use two-block partial least squares analyses and Ornstein-Uhlenbeck models of adaptive trait evolution to explore the phenotypic evolution of hard and soft cranial tissues and test hypotheses of correlated trait evolution. In pairwise analyses we found that all traits (endocast shape, neurocranium shape, rostrum shape, jaw muscle shape and residual orbit diameter) are significantly correlated. We found strongest support for a modular hypothesis of trait evolution across the whole head, with the rostrum and jaw muscles forming one module and brain, neurocranium, and eye forming the other. Within modules, traits are tightly integrated, but integration is relaxed between modules, allowing them to develop and evolve with a degree of independence. Together, these results highlight the integrated nature of the avian head and reveal that rather than being driven overwhelmingly by selection on a single trait, the shape of the avian head is a result of multiple interactions among hard and soft tissue traits.

## Introduction

Among the many phenotypic changes associated with the origin of birds, one of the most striking is the increase in brain volume relative to body size compared to non-avian theropod dinosaurs. This increase in relative brain size is also associated with differential expansion and restructuring of regions of the brain associated with cognition and vision (Balanoff et al., 2013; Early et al., 2020a; Field et al., 2025; Marugán-Lobón et al., 2016; Torres et al., 2021; Tsuboi et al., 2018; Wylie et al., 2015). In modern birds, it is clear that variation in brain structure and function underlie a broad range of complex behaviours, including sociality, vocal learning, parental care, and flight (Corfield et al., 2012; Early et al., 2020a; Güntürkün et al., 2023; Heldstab et al., 2022; Jerison, 1985; Olkowicz et al., 2016). Indeed, many researchers have proposed that evolution of brain shape and size is a key factor influencing the ecological diversification of the clade (Balanoff et al., 2016; Iwaniuk and Hurd 2005; Vincze et al., 2015). As such, understanding the factors that influence neuroanatomical evolution across the avian tree of life has been a key area of research focus (Balanoff et al., 2016; Ksepka et al., 2020; Watanabe et al., 2021).

There is strong comparative and experimental evidence that the shape and relative proportions of functional regions of the brain reflect their processing capacities because of the corresponding increase in the number of neurons (Jerison, 1973; Wylie et al., 2015). Sometimes called the “principle of proper mass,” this structure-to-function mapping is typically thought of as the primary driver of neuroanatomical evolution and allows for cognitive, sensory, and behavioural traits to be reconstructed in extinct taxa (Ashwell and Scofield, 2008; Balanoff et al., 2013 & 2016; Early et al., 2020a & b; Gold et al., 2016; Iwaniuk and Nelson, 2002; Torres et al., 2021; Watanabe et al., 2021). For example, long distance migratory birds have enlarged optic lobes, presumably as a result of selection on high visual acuity (Vincze et al., 2015). Similarly, the high cognitive ability in famously intelligent birds such as parrots and crows is reflected in the presence of a relatively larger cerebrum (forebrain) (Emery, 2006; Güntürkün et al., 2024; Iwaniuk and Hurd, 2005; Olkowicz et al., 2016).

To date, most research into the evolution of brain shape variation across birds has focused on the influence of the function of the brain itself (Balanoff et al., 2016; Güntürkün et al., 2023; Isler and van Schaik, 2006; Iwaniuk and Hurd, 2005; Olkowicz et al., 2016; Wylie et al., 2015) or of allometry (Early et al., 2020a; Kawabe et al., 2013; Marugán-Lobón et al., 2016; Torres et al., 2021; Tsuboi et al., 2018; Watanabe et al., 2021). However, the function and ontogenetic growth of the brain is strongly correlated and dependent on many other cranial structures, and these correlations might impose trade-offs or constraints that also contribute to patterns of brain shape evolution. For example, eye and brain shape are significantly correlated in birds, and differences in brain shape are further associated with orientation of the neck and thus head posture (Kawabe et al., 2013). Increased encephalisation may also impact the evolution of other soft tissues. The rostrocaudal orientation of jaw muscles in neognath birds differs markedly from the more vertically orientated jaw muscles in their extinct theropod relatives, and it has been proposed these radical changes have been caused by the lateral expansion of the brain in modern birds (Wilken et al., 2025). This ultimately led to the evolution of powered cranial kinesis in neognaths, in part to compensate for the resultant reduction in mechanical efficiency of the restructured jaw musculature. Similarly, the shape of the brain and neurocranium are tightly linked during development (Abzhanov et al., 2004 & 2006; Bhullar et al., 2015; Conith et al., 2023; Hu and Marcucio, 2009; Hu et al., 2015; Hüppi et al., 2021). This is because the brain originates early in embryological development and influences the formation of the surrounding protective skull tissue (Francis-West et al., 2003; Hu et al., 2015; Marcucio et al., 2005). As a result, changes in brain and skull shape track each other closely through both development and evolution (Fabbri et al., 2017; Watanabe et al., 2018). These lines of evidence give support for the ‘Hand in Glove’ hypothesis which proposes that the brain is the primary influence on neurocranium morphology. The neurocranium in turn provides structure and support for the surrounding soft tissues, including the jaw muscles and eye (Holliday, 2009; Kawabe et al., 2013; Richtsmeier and Flaherty, 2013). Conversely, there is evidence that development of the skull may itself affect brain shape and size. Research into craniosynostosis (premature fusion of cranial sutures) in humans has revealed that resultant changes in relative growth of the overlying skull bones can impact the shape of the enclosed brain (Richtsmeier et al., 2006; Richtsmeier and Flaherty 2013). This implies that postnatal skeletal growth may influence the subsequent development of the brain, despite the morphogenetic primacy of the brain in early embryogenesis.

The ‘Spatial Packing Hypothesis’ (SPH) predicts that the head has a finite capacity to accommodate and maintain the functional integrity of a range of structures, and that the size, shape and position of anatomical structures necessarily impact, and are impacted by, the growth and development of the surrounding tissues (Jeffery et al., 2021). One example of spatial packing trade-offs in vertebrates comes from experiments on mice with myostatin deficiency which results in increased muscle volume, including of the jaw muscles that are attached to the temporal region of the skull. Consequently, these mutant lines developed smaller brain volume than wild-type mice, suggesting that jaw muscles and the brain compete for space in the head (Jeffery and Mendias, 2014; Jeffery et al., 2021). Under normal muscle growth, however, trade-offs may not operate in the same way. Jaw muscle growth in domestic chickens (Gallus gallus) exhibits positive allometry relative to eye and brain volume, indicating that space within the head is not a limiting factor for soft tissue growth in this species (Cerio et al., 2023). In another example from birds, the brain and eyes both occupy a large volume of the head and therefore compete for space within it, with the size of the head itself defining the upper volume limit of these components (Burton, 2008). One notable result of this is that an increase in relative brain size combined with a shortened basicranial length may lead to increased basicranial flexion, driven by the restructuring of cranial traits to accommodate this increase (Lieberman et al., 2008; Marugán-Lobón et al., 2022; Ross and Ravosa, 1993). Either a trade-off in volume or a change in the shape of these structures to accommodate them both within the head may therefore be expected when any major changes in size or shape occur, but the extent to which these patterns extend to shape, rather than size, and across taxa has not been explored.

Finally, the ‘Functional Matrix Hypothesis’ states that the form of the skull, along with its growth and maintenance, are related to the functional demands of the soft tissues that are associated with the skull (Moss and Young, 1960; Moss 1971). Under this hypothesis, skull growth is mediated by the mechanical forces imposed by the non-skeletal tissues of the head, such as the growing brain and cranial muscles (Moss, 1997). The head thus develops as part of a functional complex of hard and soft tissues that is shaped by mechanical and functional demands. This hypothesis has received support from embryological experiments in mice that showed neuronal function and peripheral innervation were crucial for normal craniofacial development (Kyrkanides et al., 2011).

Together, these interrelated hypotheses leverage evidence from embryology to explain the correlated development of the skull, brain, eye and cranial muscle tissues. However, the consequences of these developmental interactions between cranial tissues are poorly understood across macroevolutionary timescales, and the effects of selection on different structures of the skull cannot be fully understood without first unravelling these interactions (Klingenberg and Marugán-Lobón, 2013). Moreover, it remains unknown how the phenotypic evolution of the brain is influenced by interactions with the other structures within the head, or if brain shape itself places constraints on the morphological evolution of these other structures. Here, we use evolutionary modelling to explore three-dimensional (3D) shape variation among the skull, brain, eye and jaw musculature across a sample of 322 extant and recently extinct avian taxa to determine how interactions among cranial traits mediate phenotypic evolution under natural selection in birds. Integrating 3D shape analysis with these methods improves upon linear and volumetric analyses by incorporating more information not only about how structures vary between species, but how their phenotypic variation impacts that of other associated traits, allowing us to describe the effects of correlated trait evolution. We first investigate pairwise evolutionary correlations between hard and soft tissue structures using a two-block partial least square (PLS) approach. We then compare several competing evolutionary hypotheses by integrating these traits across the head for the first time with a novel approach using Ornstein-Uhlenbeck (OU) models of adaptive trait evolution, implemented in *mvSLOUCH* (Bartoszek et al., 2012 & 2024) to determine the best supported model for coordinated trait evolution. We expect that trade-offs in shape among hard and soft tissue components of the avian head play an important role in defining the range of morphologies seen in crown birds, and that these trade-offs will mirror those proposed in shaping the developing head through embryology and ontogeny. Ultimately, we predict that the brain plays a key role in influencing the phenotypic evolution of the avian head, but that its effects are moderated by evolutionary interactions between the surrounding traits of the head.

## Methods

### Mesh generation

We assembled a dataset of CT scans of the skulls of 322 birds (311 extant, 11 extinct; see Supplementary Information for specimen sources). Taxa were chosen to represent the taxonomic and morphological diversity of modern birds, incorporating 43 of 44 Orders, 188 of 254 Families, and 322 of 2392 Genera (Fig. 1; Gill et al., 2024). We created three-dimensional (3D) skull meshes from µCT data (segmented in Dragonfly 2022.2 and VGStudio MAX 3.5.x) and decimated them to 1 million faces. We used the ‘endomaker’ function in the R package *Arothron* (Profico et al., 2023) to create endocasts of all specimens. We manually removed the remnants of cranial nerve (CN) V from the endocast, which superficially overlays the optic lobe, to allow for consistent surface semilandmark placement on this structure.

**Figure 1:**
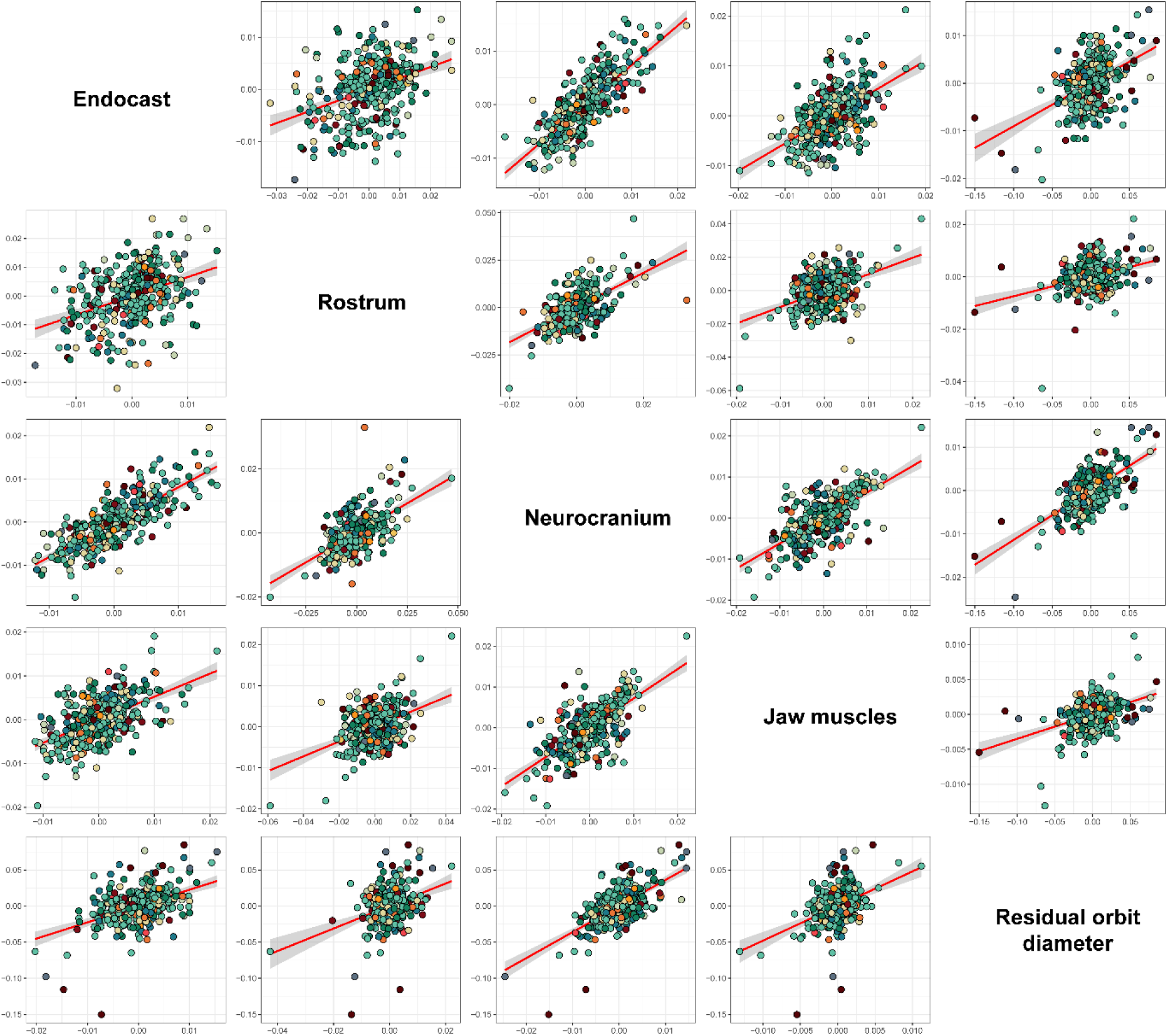
Results of 2-block PLS analyses for all traits. Best fit lines are shown in red for each plot, with 95% confidence intervals represented by shaded area. Each trait is read as the y-axis (horizontally) or x-axis (vertically) from its position on the diagonal.

### Phylogeny

We generated an informal supertree containing all 11 extinct and 311 extant taxa in our sample. Recent phylogenetic analyses using genomic data have generated new hypotheses regarding the broad-scale (i.e., family level and higher) relationships among Neoaves (Stiller et al., 2024; Supplementary Fig. S1). We thus used this recent topology as a backbone tree upon which to graft sub-family-level relationships derived from a comprehensive clade-wide phylogeny (Jetz et al., 2012). We downloaded 1,000 trees from posterior distribution of species-level Neoaves phylogenies from BirdTree.org using the “Hackett Stage 2” topologies (Jetz et al., 2012). Using Tree Annotator, we generated a single maximum clade credibility MCC tree from this posterior sample under the “target heights method” (Drummond and Rambaut, 2007). We extracted individual clades from this MCC and grafted them to the corresponding positions in the backbone tree using established approaches (Cooney and Thomas, 2021). The resulting supertree contains 9993 extant taxa with the broad-scale phylogenetic relationships and divergence times supported by the recent genomic analysis.

Finally, we added extinct taxa that are not present in the BirdTree.org (Jetz et al., 2012) topology using two approaches. For very recently extinct taxa with well-resolved phylogenetic affinities (e.g*., Ninox albifacies, Moho braccatus*) we simply substituted them with their most closely related extant taxa (*Ninox rufa, Hypocolius ampelinus*, respectively; Oliveros et al., 2019). For other taxa that are more distantly related from extant taxa (i.e., *Aptornis otidiformis*, *Cnemiornis calcitrans, Diaphorapteryx hawkinsi*, *Emeus crassus*, *Eudyptes warhami*, *Pezophaps solitaria*, *Raphus cucullatus*) we used the ‘tree.merger’ function of the *RRphylo* R package (Castiglione et al., 2018) to attach extinct taxa to known positions on the topology (Cole et al., 2019; Dumbacher, 2003; Garcia-R et al., 2014; Musser and Cracraft, 2019; Shapiro et al., 2002; Yonezawa et al., 2017).

### Geometric morphometrics

We applied landmarks to the right hemisphere of both the skull and endocast using templates adapted from Mitchell et al. (2021) and Watanabe et al. (2021) respectively, using Stratovan Checkpoint v. 2020.10.13.0859. Discrete anatomical (Bookstein Type I and II) landmarks were used to describe homologous points, and semilandmarks curves were used to capture homologous curves and sutures (Supplementary Information). The skull landmark template was modified to more comprehensively quantify jaw muscle shape (Adductor Mandibulae Externus and Depressor Mandibulae; Baumel et al., 1993) by placing semilandmark curves on the skull along the margins of the fossa musculorum temporalium and impressio musculi depressor mandibulae as proxies for muscle shape (Livezey et al., 2006; Supplementary Figures S1 and S2). This resulted in a total of 20 anatomical and 190 semilandmarks in 21 curves for the skull, and 14 anatomical and 72 semilandmarks in 15 curves for the endocast (Supplementary Information). The homologous structures of the endocast (cerebrum, optic lobe, cerebellum and medulla) have well-defined borders but the globular, smooth surfaces, have no reliably identifiable points for placing landmarks. Large areas of the endocast surface therefore cannot be captured using anatomical landmarks or semilandmark curves. Following Watanabe et al. (2021) we used a semi-automated method to place surface semilandmarks on the surface of these structures. We used a single specimen to create a template of 103 surface semilandmarks and then used a semi-automated procedure in the R package *Morpho* (Schlager, 2017) to apply the patch to all endocast specimens (Bardua et al., 2019).

We slid semilandmarks to minimise bending energy and then reflected landmarks across the midline for both skull and endocast to ensure bilateral symmetry, a protocol which prevents lateral deviation from the midline that can be induced during landmark alignment (Zelditch et al., 2004). Symmetrised landmark data were aligned with a general Procrustes alignment (GPA) using the ‘procSym’ function in the R package *Morpho* (Schlager, 2017). We performed global alignment separately for the whole skull and endocast landmark set, as well as for landmark subsets of the skull (neurocranium, rostrum, and jaw muscles). After alignment we removed reflected landmarks and performed all subsequent analyses on the right-side landmarks for each landmark subset.

We explored shape variation by performing a principal components analysis (PCA) on the Procrustes-aligned shape data for the whole skull and for each subset of landmarks (endocast, neurocranium, jaw muscles and beak), using the ‘gm.prcomp’ function in the R package *geomorph* (Adams et al., 2023).

### Residual orbit diameter

To account for the non-isometric scaling of eye size with body mass (Burton, 2008) we adapted the formula for calculating encephalization quotient (EQ) to calculate residual orbit diameter (ES) for each species with the formulae *ES = ES_a_/ES_e_* (Jerison, 1973; Roth and Dicke, 2005). We measured observed orbit diameter (*ES_a_*) from the maximum linear diameter of the orbit for each specimen. We then performed a linear regression of log-transformed observed orbit diameter against log-transformed body mass and used the fitted values from this regression (i.e., *ES_a_* – *residual*) to obtain expected orbit diameter (*ES_e_*) for each taxon. We took mean body mass for each species from AVONET (Tobias et al., 2022) for extant birds, and used a combination of published values and scaling equations from Field et al. (2013) to estimate body mass from hindlimb bone dimensions for extinct taxa (see Supplementary Table S1 for details).

### Pairwise correlation

To test evolutionary correlations between traits, we performed two-block partial least squares (PLS) analyses for each pairwise trait combination (beak shape, endocast shape, neurocranium shape, jaw muscle shape and ES) using the ‘phylo.integration’ function in the R package *geomorph* (Adams and Felice, 2014; Adams and Collyer, 2016; Baken et al., 2021; Adams et al., 2023) with Procrustes-aligned landmark data, and log-transformed values for ES.

### Evolutionary modelling

We modelled the correlated evolution of cranial hard and soft tissues to test the hypothesis that selection for brain shape influences the evolution of the skull, eyes, and jaw muscles, and vice versa. We used a multivariate Ornstein-Uhlenbeck (OU) model of adaptive evolution as implemented in the R package *mvSLOUCH* v2.7.6 (Bartoszek et al., 2012 & 2024). Under the multivariate OU model, traits are pulled toward an adaptive optimum or optimum by an attractive force that is interpreted as natural selection and strength of this selection is described by the matrix **A**. The diagonal elements of the **A** matrix describe the selection on individual traits and the off-diagonal elements represent the degree to which traits’ paths toward the optimum influence each other (Bartoszek et al., 2012 & 2024). The strength of the mvSLOUCH approach to modelling multivariate OU processes is that specific hypotheses of trait interactions and trade-offs can be evaluated by specifying the properties of the **A** matrix.

We formulated twelve hypotheses to investigate evolutionary trade-offs among our sampled traits (Table 1), constructing an **A** matrix for each hypothesis to define trait interactions (Bartoszek et al., 2024). Our models were based on the three main hypotheses of correlated trait evolution (Hand in Glove, Spatial Packing, and Functional Matrix). For each hypothesis we designed a model of trait interactions which best fits the set of traits that we sampled and designed a second model that involves feedback between traits. We designed a third Spatial Packing hypothesis in which jaw muscle shape is not directly influenced by brain shape but instead evolves in response to rostrum shape. This is based on the observation that there is minimal trade-off between brain size and jaw muscle size during embryonic and early postnatal growth of *Gallus gallus* (Cerio et al., 2023). We also included ‘Ecological Selection’ models, which imply that beak shape, which dominates morphological variation across birds (Bright et al., 2016; Felice et al., 2019), is a direct result of selection as opposed to the developmental byproduct of shape changes in other traits. In these models the shape of other traits within the head are a result of shared developmental pathways and are consequently influenced by selection on beak shape. Finally, we created a ‘Modular’ model, in which the head is divided into two non-interacting phenotypic modules, the neurosensory module, composed of the endocast, neurocranium and residual orbit diameter, and the feeding module, composed of the rostrum and jaw muscles. These regions comprise the two major components of the kinetic avian skull (Wilken et al., 2025) and are derived from separate developmental centres, namely the mandibular cranial neural crest (rostrum) and mesoderm (neurocranium; Bhullar et al., 2015). This model was included to account for the possibility that this developmental modularity leads to mosaic evolution in the avian head (Felice and Goswami, 2018) and may be an important factor in the evolution of cranial kinesis in modern birds (Wilken et al., 2025). All hypotheses were contrasted with the Brownian motion (BM) null model of no selection on any trait. We conducted two sets of analyses for each model to explore differences in trait evolution that are not connected with adaptation, e.g. developmental correlations. In the first set we assigned the stochastic perturbations matrix (**Σ**_yy_) to be diagonal, implying no interactions between the traits in the non-adaptive component of evolution. In the second set, **Σ**_yy_ was upper-triangular, indicating interacting evolution of the traits due to non-adaptive effects.

**Table 1:**
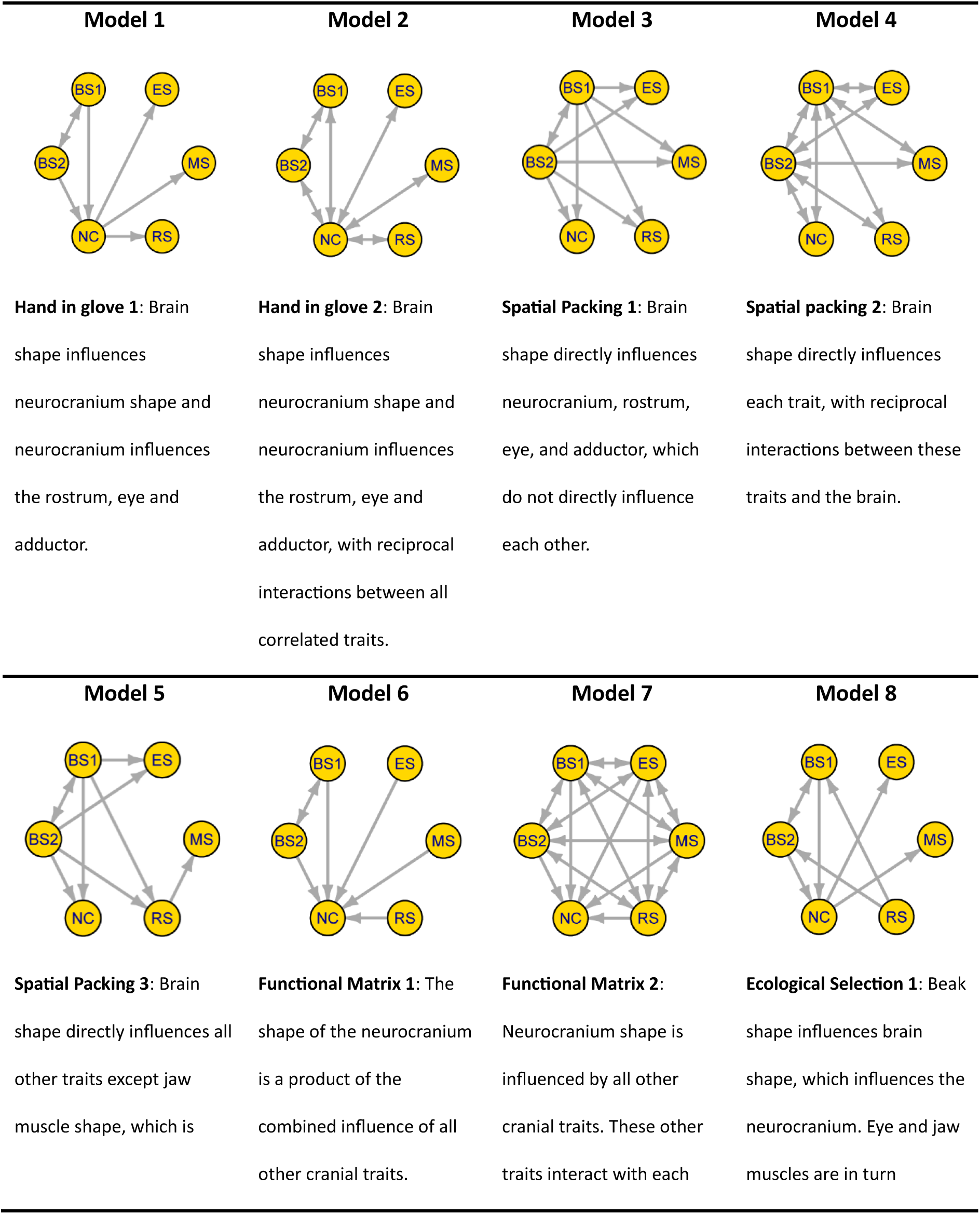
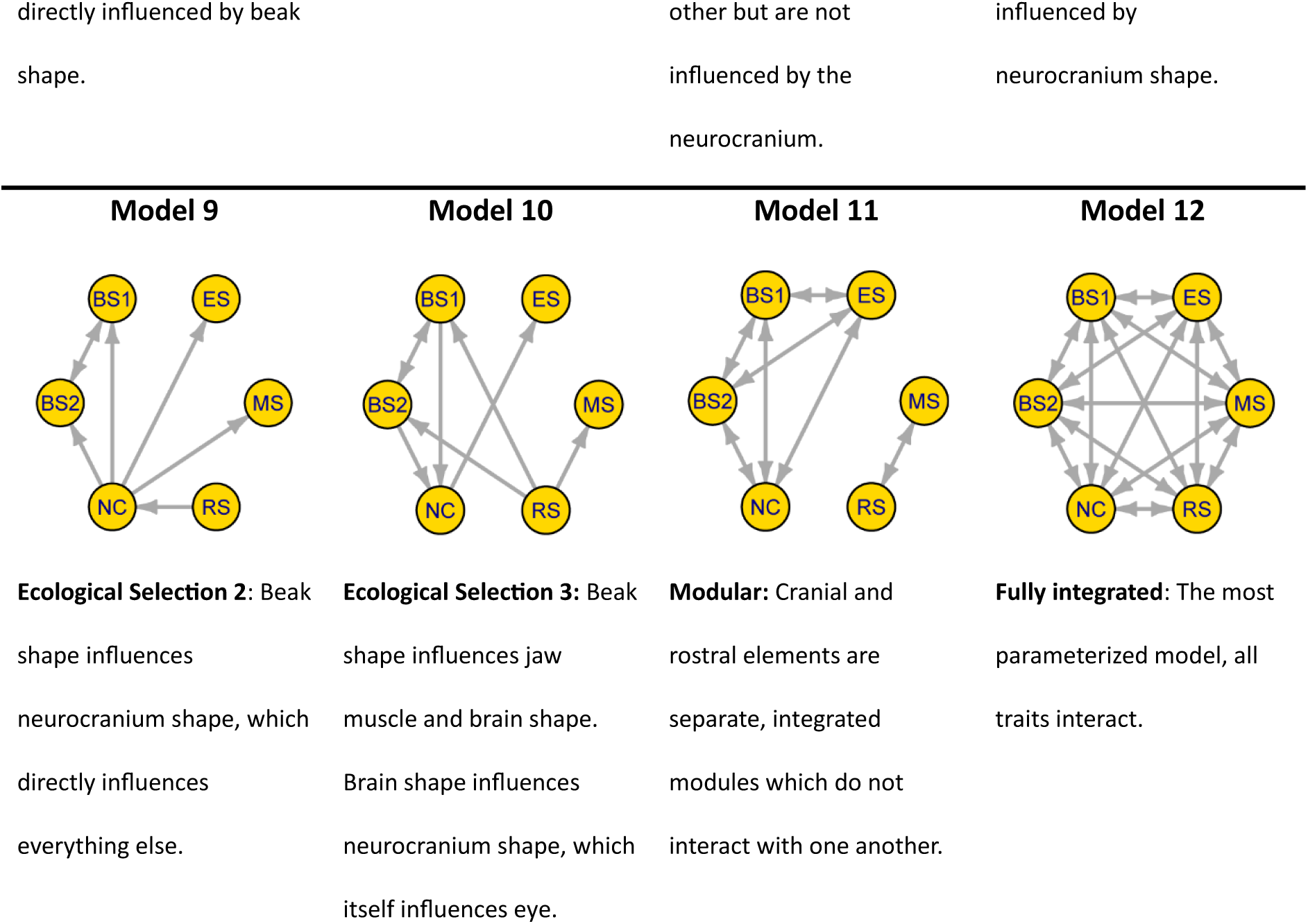
Correlated trait models tested with *mvSLOUCH*. Diagrams at left represent model interactions. Circles represent traits and arrows represent the direction of trait influence. Traits are NC = neurocranium shape; BS1 and BS2 = PCs 1 and 2 of endocast shape, respectively; ES = residual orbit diameter; MS = jaw muscle shape; RS = rostrum shape. See Supplementary Table S3 for corresponding **A**-matrices.

We compared Akaike information criterion (AICc) scores to determine the best-supported model. We determined confidence intervals for each model parameter by performing 100 parametric bootstraps on the highest likelihood model. The tempo of adaptation, expressed as phylogenetic half-lives for each eigenvector of the **A** matrix in the best-supported model, is also provided by *mvSLOUCH*. Half-lives represent rates of adaptation of the set of traits towards the primary optimum along the directions in trait space described by that eigenvector, measured as the time needed for the effect of the initial state to be halved (Bartoszek et al., 2012). It is important to note that, because this is a multivariate analysis, these half-lives refer to the adaptation of a group of complex traits rather than a single trait, as would be implied in a univariate model.

Previous research has suggested broader-scale phylogenetic patterns of relative brain size among extant birds (Iwaniuk and Hurd, 2005; Marugán-Lobón et al., 2016; Torres et al., 2021). Australaves is a monophyletic clade composed of Passeriformes, Psittaciformes, Falconiformes and Cariamiformes, and contains taxa with the largest relative brain sizes among extant birds (Olkowicz et al., 2016; Güntürkün et al., 2023). Palaeognathae, by contrast, generally have low relative brain sizes compared to most birds (Ashwell and Schofield, 2008) and do not possess the highly kinetic craniofacial and palatal region of neognaths (Wilken et al., 2025). We therefore subdivided our dataset into Australaves, Palaeognathae and ‘other’ to investigate how trait optima differ between these groups in the best supported model.

## Results

Variation in whole-skull shape in the dataset is dominated by variation in beak shape (Supplementary Figure S4). The major axis of shape variation in whole-skull shape data (PC1) is elongation, representing 45% of total skull shape variation, and with short-skulled (brachycephalic) taxa at negative PC values and long-skulled (dolichocephalic) taxa at positive PC values. This is followed by variation in beak depth (PC2; 10.3% of total shape variation) and relative orientation of beak and foramen magnum (PC3; 9.86% of total shape variation). Similar patterns are seen when the skull is broken down into component parts. The major axis of shape variation in the rostrum is elongation (PC1; 46.1% of total shape variation), as it is with neurocranium shape (PC1; 25.2% of total shape variation, Supplementary Figure S6). Disparity in jaw muscle shape is largely driven by differences in the relative size of the adductor mandibulae and the depressor mandibulae, with 48% of shape variation accounted for by relative expansion of the adductor mandibulae (Supplementary Figure S7). The first two PC axes of the endocast make similar contributions to total shape variation (PC1 = 33.1%; PC2 = 25.6%) and correspond to relative expansion of the cerebrum and flexion of the basicranial angle, and elongation of the whole brain, respectively (Supplementary Fig. S8).

All trait pairs are significantly correlated (p < 0.05), indicating strong evolutionary integration in the head of extant birds (Fig. 2; Supplementary Table S1). Elongation is the dominant form of shape variation of individual traits in each pairwise analysis of trait correlation (Supplementary Figure S9), usually corresponding to the shape variation seen in the first principal component of each trait (Supplementary Figures S4 to S8). This effect is most obvious with rostrum shape, which shows clear elongation in correlation with neurocranium elongation (Supplementary Figure S9F and S9J), and with endocast elongation and ventral flexion (Supplementary Figure S9E and S9A). Similarly, lengthening of the rostrum is correlated with enlargement and elongation of the attachment area of the adductor mandibulae (Supplementary Figure S9G and S9N). The dominant form of neurocranium covariation with all other traits is also elongation. For example, more elongate neurocrania are associated with a relatively enlarged adductor mandibulae (Supplementary Figure S9K and S9O). Basicranial flexion of the endocast is apparent to varying amounts in all two-block PLS analyses. This effect is most obvious with residual orbit diameter, where higher flexion angle is associated with relatively larger orbits (Supplementary Figure S9D). Flexion of the brain in these instances resembles the shape variation seen along PC1 of the PCA for brain shape, where more acute angles are associated with larger cerebrum (Supplementary Figure S8).

The mvSLOUCH analysis recovered best support for the Modular model of trait interactions with diagonal **Σ**_yy_ with an AIC weight of 1 (Table 2). In this model, traits are divided into two integrated modules, the neurosensory module (composed of the brain, neurocranium and residual orbit diameter) and feeding module (composed of the rostrum and jaw muscles). Selection in each trait within a module influences the evolutionary trajectory of each other trait in the module another, but there are no adaptive interactions between the two modules. Support for diagonal **Σ**_yy_ indicates that there are no interactions between traits in the noise component of evolution in this model. These interactions are typically associated with developmental constraints or covariation with other, unmeasured traits (Bartoszek et al., 2012 & 2023). The next best supported models are Integrated and Hand in Glove 2, with ΔAICc values of 32.27 and 42.77, respectively. The five best supported models all involve reciprocal interactions, suggesting that phenotypic integration plays an important role in the evolution of the avian head.

**Table 2:** Model support from mvSLOUCH analysis.

| Rank | AICc values | Model | $\Delta AICc$ | AIC weights | Relative log likelihoods |
| --- | --- | --- | --- | --- | --- |
| 1 | 1752.81 | **Modular** | 0.00 | 1 | 1 |
| 2 | 1785.08 | **Fully integrated** | 32.27 | 0 | 0 |
| 3 | 1795.58 | **Hand in Glove 2** | 42.77 | 0 | 0 |
| 4 | 1808.52 | **Spatial Packing 2** | 55.72 | 0 | 0 |
| 5 | 1859.33 | **Ecological Selection 2** | 106.52 | 0 | 0 |
| 6 | 1885.97 | **Hand in Glove 1** | 133.17 | 0 | 0 |
| 7 | 1910.38 | **Functional Matrix 1** | 157.57 | 0 | 0 |
| 8 | 1942.17 | **Spatial Packing 1** | 189.36 | 0 | 0 |
| 9 | 1953.39 | **Spatial Packing 3** | 200.58 | 0 | 0 |
| 10 | 1990.74 | **Ecological Selection 3** | 237.93 | 0 | 0 |
| 11 | 1999.21 | **Ecological Selection 1** | 246.40 | 0 | 0 |
| 12 | 2058.633 | **Functional Matrix 2** | 305.82 | 0 | 0 |

Rates of adaptation, measured as phylogenetic half-lives, are relatively consistent across eigenvectors and range from 60.23% to 76.57% of tree height, translating to 68.1 to 86.6 Ma (Table 3). Each eigenvector reveals the adaptive contribution of traits via loadings. For example, eigenvector 2 (e_2_) is driven by residual orbit diameter (ES; 0.82) and the first component of brain shape (BS1; 0.51) but the second component of brain shape (BS2; -0.14) and neurocranium shape (NC; -0.21) have negative eigenvalues. Therefore, adaptation toward the trait optimum for BS1 and residual eye size is correlated with evolution away from the trait optimum for neurocranium shape and BS2. The high negative loadings in e_3_ (first component of brain shape) and e_5_ (neurocranium shape) imply that these eigenvectors are dominated by the respective traits moving away from the trait optima. Because rostrum shape (RS) and jaw muscle shape (MS) do not interact with other traits in this model they only contribute to e_1_ and e_6_. MS contributes the most to e_1_ (1.00) and RS to e_6_ (1.00). Jaw muscle shape is effectively the fastest evolving trait because it dominates e_1_ (the eigenvector with the highest rate of adaptation). This may be because jaw muscle shape has high phenotypic plasticity and is more readily able to adapt to new optima, but it is worth noting that the range of adaptation rates across all eigenvectors is low, meaning that each eigenvector of selection has similar strength.

**Table 3:** Phylogenetic half-lives with 95% parametric bootstrap (100 bootstrap replicates). Eigenvectors (columns) are arranged L-R in descending order of adaptive rate, represented by half life as a percentage of total tree height (bottom row). Abbreviations in other rows correspond to traits: Jaw muscle shape (MS), residual orbit diameter (ES), endocast shape (BS1 and BS2), neurocranium shape (NC) and rostrum shape (RS).

| Directions (eigenvectors) |  |  |  |  |  |  |
| --- | --- | --- | --- | --- | --- | --- |
|  | e<sub>1</sub> | e<sub>2</sub> | e<sub>3</sub> | e<sub>4</sub> | e<sub>5</sub> | e<sub>6</sub> |
| **MS** | 0.999744 | 0.000000 | 0.000000 | 0.000000 | 0.000000 | 0.068764 |
| **ES** | 0.000000 | 0.820496 | 0.146992 | -0.014438 | 0.054588 | 0.000000 |
| **BS1** | 0.000000 | 0.510677 | -0.957219 | 0.023156 | -0.119173 | 0.000000 |
| **BS2** | 0.000000 | -0.141449 | 0.047982 | 0.981254 | -0.041832 | 0.000000 |
| **NC** | 0.000000 | -0.214448 | 0.244589 | 0.190775 | -0.990489 | 0.000000 |
| **RS** | -0.022632 | 0.000000 | 0.000000 | 0.000000 | 0.000000 | 0.997633 |
| **Half-life** | 60.23% | 65.51% | 65.81% | 66.99% | 68.82% | 76.57% |

Trait optima values vary between subsets of taxa for all traits and is especially noticeable in brain shape (Fig. 3). Palaeognathae and Australaves have positive optima for the first axis of brain shape variation (BS1), indicating that these groups have tended to evolve a relatively larger cerebrum than taxa in the ‘other’ group, which have a negative shape optimum value. In contrast, Palaeognathae have a negative optimum value along the second axis of brain shape variation (BS2), corresponding to a more elongate endocasts with a posteriorly oriented brain stem. Palaeognathae also differ noticeably from other groups in neurocranium shape (NS), having an elongate shape optimum, and residual orbit diameter (ES), having a relatively smaller orbit diameter. All three groups have a similar optimum for rostrum shape (RS) and jaw muscle shape (MS), which in both instances broadly correspond to the mean value for these traits.

**Figure 3:**
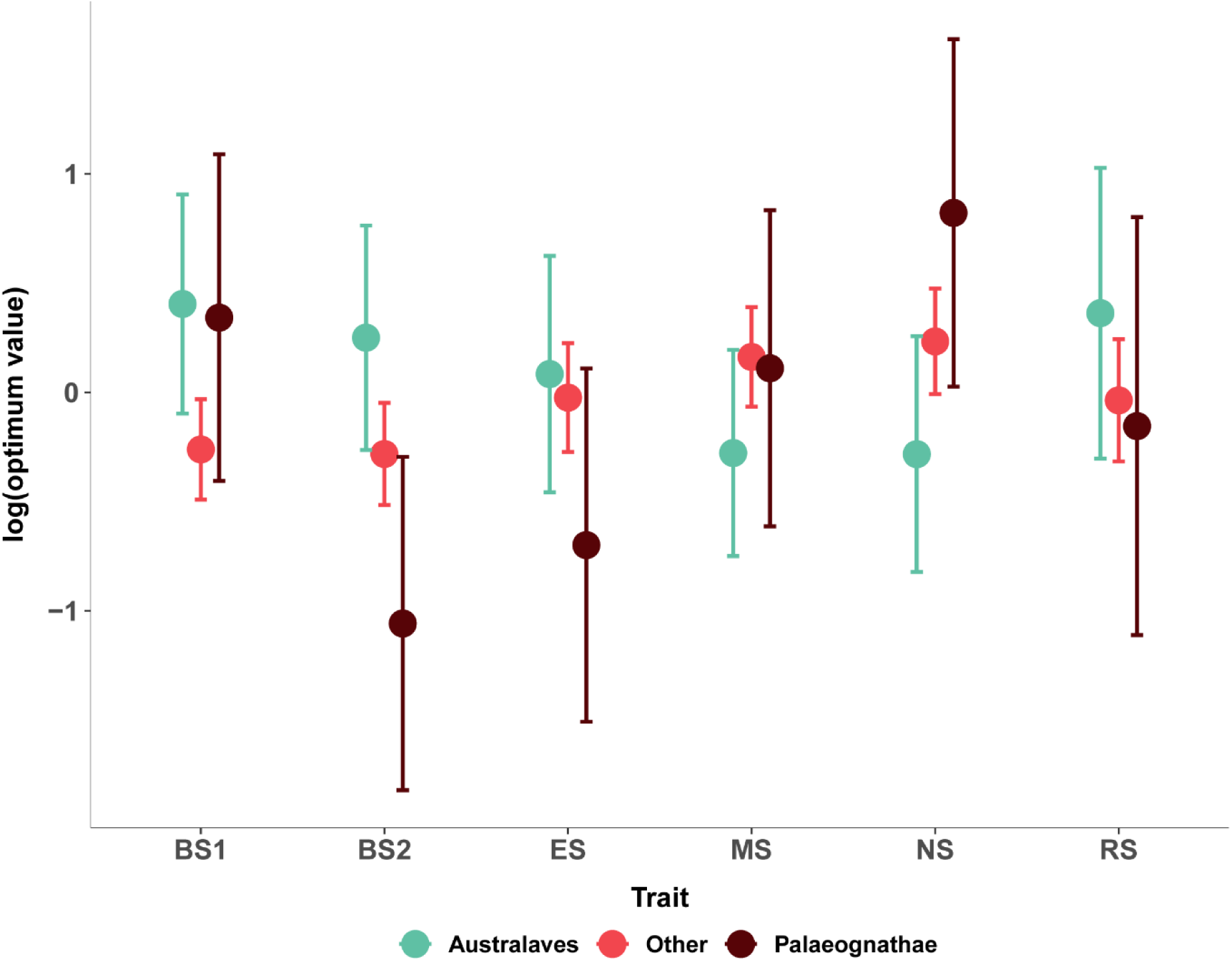
Deterministic optimum values of all traits for three subsets of species. Values are grouped according to trait (x-axis) and coloured according to group.

## Discussion

Our findings reveal that the avian head is shaped by the coordinated coevolution of skeletal and soft tissues, and that the interaction of these traits play a major role in both generating and constraining phenotypic diversity in Aves. In addition to known correlations between brain and neurocranium shape (Fabbri et al., 2017; Marugán-Lobón et al., 2022) and between brain shape and eye size (Kawabe et al., 2013), our analyses reveal significant pairwise correlations between all examined traits (brain shape, rostrum shape, neurocranium shape, jaw muscle shape and residual orbit diameter). A common pattern seen in the two-block PLS analyses is that each trait yields similar trends of shape variation in response to shape variation in other traits. For example, rostrum shape varies along a continuum from short and curved to elongate and straight in all pairwise analyses (Supplementary Figure S9). Similarly, the degree of ventral flexion of the endocast is associated with specific morphologies in all other traits (Supplementary Figure S9). These results suggest that patterns of covariation of cranial traits are broadly consistent in birds and implies that trade-offs in shape between traits play an important role in avian cranial evolution.

The best supported model describes an adaptive evolutionary process with a modular pattern of trait interactions where integration between the feeding and neurosensory modules is relaxed, allowing some evolutionary independence (Table 2). This may be because these two modules are largely derived from different developmental tissues, the mandibular cranial neural crest (rostrum) and mesoderm (neurocranium). Relaxed developmental separation between elements can promote mosaic evolution by allowing them to adjust development independently in response to selection (Felice and Goswami, 2018; Maddin et al., 2016), despite the shared signalling pathways which influence both brain and skull development in vertebrates, including FGF, TGFβ, Wnt, Hedgehog and Notch (Richtsmeier and Flaherty, 2013; Bhullar et al., 2015; Neben and Merrill, 2015). Changes in the developmental timing and structural arrangement of the head have been linked to the early evolution of crown birds and the evolution of the avian beak and palate within Archosauria (Bhullar et al., 2012 & 2016; Hüppi et al., 2021; Plateau and Foth, 2020; Yang and Sander, 2018). In more closely related taxa, studies of Darwin’s finches (Fringillidae, Passeriformes) have shown that beak elongation in some taxa is due to higher levels of calmodulin (CaM) expression in the developing rostrum (Abzhanov et al., 2006). Similarly, misexpression of Bmp4 in the mesenchyme of the upper rostrum in embryonic domestic chickens results in the development of deep beak morphology resembling that of the Galapagos finch *Geospiza magnirostris* (Abzhanov et al., 2004). The ability of regions of the head to develop and evolve with this degree of independence is a hallmark of modularity (Zelditch and Goswami, 2021) and has allowed beak shape to dominate morphological variation in adult birds (Supplementary Figure S4; Bright et al., 2016; Felice et al., 2019). Beak shape has been linked to foraging ecology, suggesting it is under strong adaptive selection (Bright et al., 2019; Natale and Slater, 2022; Navalón et al., 2019). Moreover, and the evolution of powered kinesis in the neognath skull has been suggested as a key evolutionary innovation, increasing mechanical efficiency of jaw muscles and improving craniofacial dexterity in the absence of dextrous forelimbs (Wilken et al., 2025). Relaxed integration between the feeding and neurosensory modules gives the beak and jaw muscles greater freedom to explore novel morphologies and adapt their mechanical properties to adapt to a range of food sources without severely impacting brain shape, which may be responding to different adaptive pressures or functional needs.

Our findings do not contradict any of the developmental hypotheses outlined in the introduction (Hand in Glove, Spatial Packing and Functional Matrix), and there is experimental evidence that each plays a role in shaping the vertebrate head during embryogenesis and development (Kyrkanides et al., 2011; Marugán-Lobón and Buscalioni, 2006; Marugán-Lobón et al., 2022; Richtsmeir and Flaherty, 2013). Notably, the key role of the brain in morphogenesis of the vertebrate head is known from many developmental studies (Douarin, 2004; Hanken and Thorogood, 1993; Hu and Marcucio, 2009; Hu et al., 2015; Hüppi et al., 2021; Marcucio et al., 2005; Richtsmeier and Flaherty, 2013). Our findings confirm that that the brain is evolutionarily integrated with the skull and that reciprocal interactions between brain shape, neurocranium shape, and eye size combine to influence head morphology across crown birds. Correlation between traits suggests that important trade-offs exist between regions of the avian head (Fig. 1) and lends some support to the Spatial Packing Hypothesis. Whereas the OU models that we label Spatial Packing Hypothesis (Table 1, Models 3-5) each involve brain shape directly influencing the evolution of other traits (Marugán-Lobón et al., 2022), the best-supported model in our analysis (Table 1, Model 11) could also be interpreted as a form of spatial packing because of the high degree of reciprocal interactions among traits. Trade-offs between all traits may not be universal, however, and evidence for a trade-off between brain shape and jaw muscle morphology is more mixed. On one hand, the two block PLS result indicates that there is a correlation between these two traits. Conversely, OU models that include causal interactions between brain shape and jaw shape have much lower likelihood than those without (Table 2).

Incorporating reciprocal interactions into models generally results in better support (Table 2), suggesting that peripheral soft tissues may moderate phenotypic evolution of the neurocranium and may provide some support to the Functional Matrix Hypothesis. Nonetheless, our interpretation of this hypothesis, where variation in brain shape, eye size, jaw muscle shape and rostrum shape influence the shape of the neurocranium, is not well-supported (Table 2). Although the Functional Matrix Hypothesis plays an established role in development (Kyrkanides et al., 2011), its effects on evolutionary morphology may be masked by larger-scale trends in shape evolution.

We have shown that interactions among traits in the avian head play an important role in regulating phenotypic evolution across crown birds. The brain is known to be a key influence on the development of surrounding cranial tissues, but our results emphasise the role that other cranial elements play in moderating morphological evolution of the avian head. Our sample is composed exclusively of contemporary crown birds, and we cannot determine if the patterns of trait coevolution that we have found extend to more basal taxa that may not display the traits we associate with modern birds (Field et al., 2025). Incorporating basal taxa may thus reveal different patterns of trait evolution earlier in the evolutionary history of birds. Nonetheless, our sample is representative of the phenotypic diversity of modern birds and reveals a great deal about their >100 Ma evolutionary history (Brocklehurst and Field, 2024). The coordinated coevolution of cranial traits and the pattern by which they influence one another across Aves highlights the importance of assessing the adaptive evolution of phenotypic traits in their developmental and evolutionary context and ultimately has implications for broad-scale studies of phenotypic evolution and trait integration.

## Acknowledgements

We thank Brett Clark, Mike Day and Agnese Lanzetti (NHMUK) for their help with specimen access and scanning, and other collections staff and researchers who made scan data available on MorphoSource. Ryan Marek (UCL) and Katherine Steinfield (University of Konstanz) provided additional scans. Laura Porro (UCL) found and collected the woodcock (*Scolopax rusticola*) used in this study. This work was funded by UK Research and Innovation (UKRI) under the UK government’s Horizon Europe funding guarantee EP/Y010256/1 – BRAIN-DRAIN to RNF.

## Author contributions

**AK**: Conceptualisation, data curation, formal analysis, investigation, methodology, visualisation and writing. **TW**: Data curation, investigation and writing. **CME:** Conceptualisation and writing. **RNF**: Conceptualisation, data curation, formal analysis, investigation, methodology, project administration, visualisation and writing.

## Notes

### Competing Interest Statement

The authors have declared no competing interest.

