## Supplementary information for "Avian cranial evolution is influenced by trade-offs in shape between hard and soft tissue traits"

Andrew Knapp[1,2]*, Taylor West[1], Catherine M Early[3], Ryan N Felice[1]

1. University College, London

2. The Natural History Museum, London

3. Science Museum of Minnesota

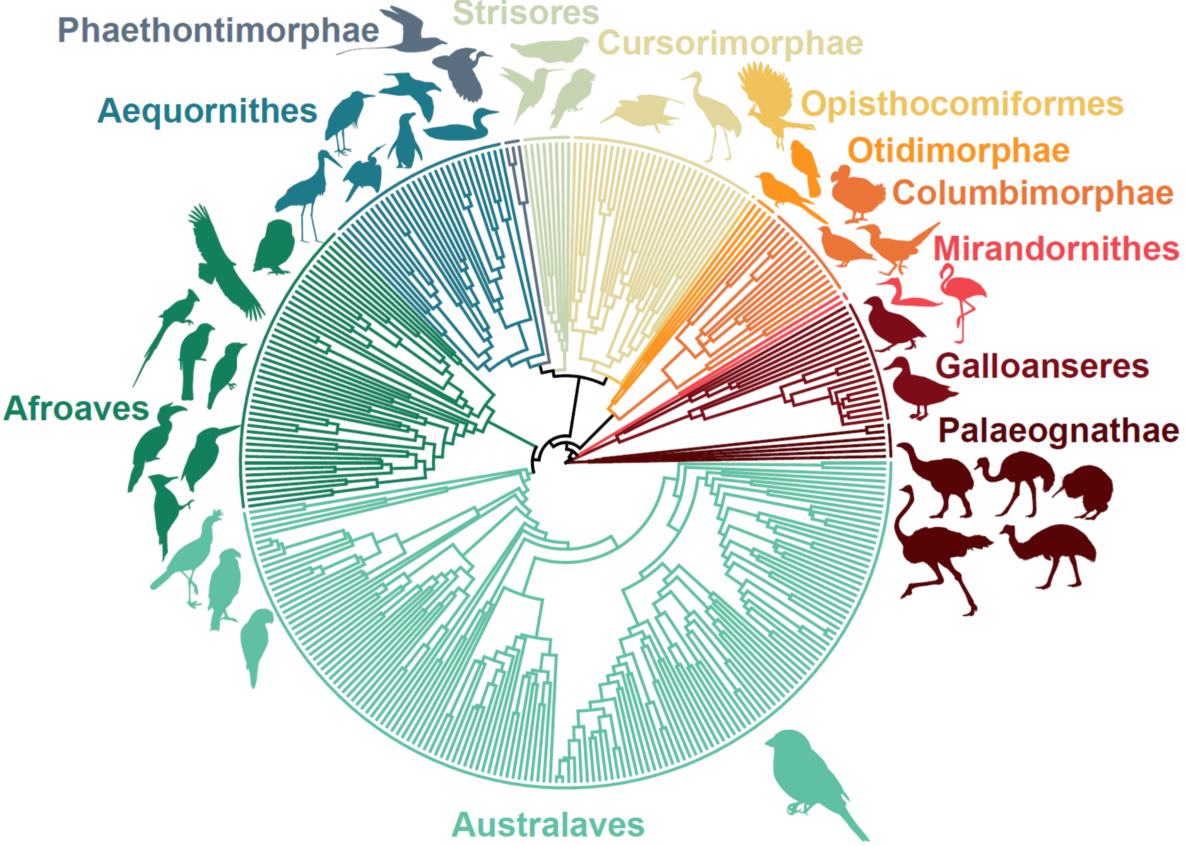


**Figure S1: Phylogeny of study sample.** Branches are coloured by major clades, defined by Stiller et al., 2024. Silhouettes represent major Orders.

**Landmark layout**

**Skull**

**Anatomical landmarks**

1. Anterior tip of rostrum
2. Posterior-most point of palatine on midline
3. Dorsal contact of jugal with maxilla
4. Dorsal contact of beak with neurocranium at craniofacial hinge, on midline
5. Contact between supraoccipital and parietal, on midline
6. Dorsalmost point of foramen magnum on midline
7. Ventralmost point of foramen magnum on midline
8. Margin of occipital condyle and basioccipital on midline
9. Contact point of basioccipital and parasphenoid on midline
10. Lateral contact of maxillary process and neurocranium at craniofacial hinge
11. Contact point of lachrymal and orbital rim
12. Distal tip of postorbital process
13. Distal tip of depressor mandibulae process
14. Ventral margin of depressor mandibulae fossa, on margin of paraoccipital process

15. Ventral-most point of margin of optic nerve foramen
16. Midline of basisphenoid level with posterior-most point of palatines
17. Ventral contact of jugal and maxilla
18. Ventral contact point between jugal and quadrate
19. Ventral-most contact point between quadrate and pterygoid
20. Anterior point of pterygoid at contact with palatine

**Semilandmark curves**

1. Lmks 1 – 2, along underside midline of premaxilla and medial margin of palatine
2. Lmks 2 – 3, along lateral margin of palatine
3. Lmks 1 – 3, along labial margin of beak
4. Lmks 3 – 4, along craniofacial hinge
5. Lmks 4 – 1, along dorsal midline of beak
6. Lmks 4 – 5, along dorsal midline of cranium
7. Lmks 5 – 6, along posterior midline of cranium
8. Lmks 6 – 7, around margin of foramen magnum
9. Lmks 7 – 8, around base of occipital condyle
10. Lmks 8 – 9, along ventral midline of basioccipital
11. Lmks 9 – 5, along nuchal crest
12. Lmks 10 – 11, along frontal – lachrymal suture
13. Lmks 11 – 12, around orbit margin
14. Lmks 12 – 13, along temporal crest delineating temporal fossa
15. Lmks 13 – 14, along margin of depressor mandibulae fossa
16. Lmks 14 – 16, along lateral margin of parasphenoid and basisphenoid
17. Lmks 16 – 9, along midline of basisphenoid and basioccipital
18. Lmks 17 – 18, along lateral margin of jugal bar
19. Lmks 18 – 19, along posterior margin of articular surface of quadrate
20. Lmks 19 – 18, along anterior margin of articular surface of quadrate
21. Lmks 19 – 20, along lateral margin of pterygoid

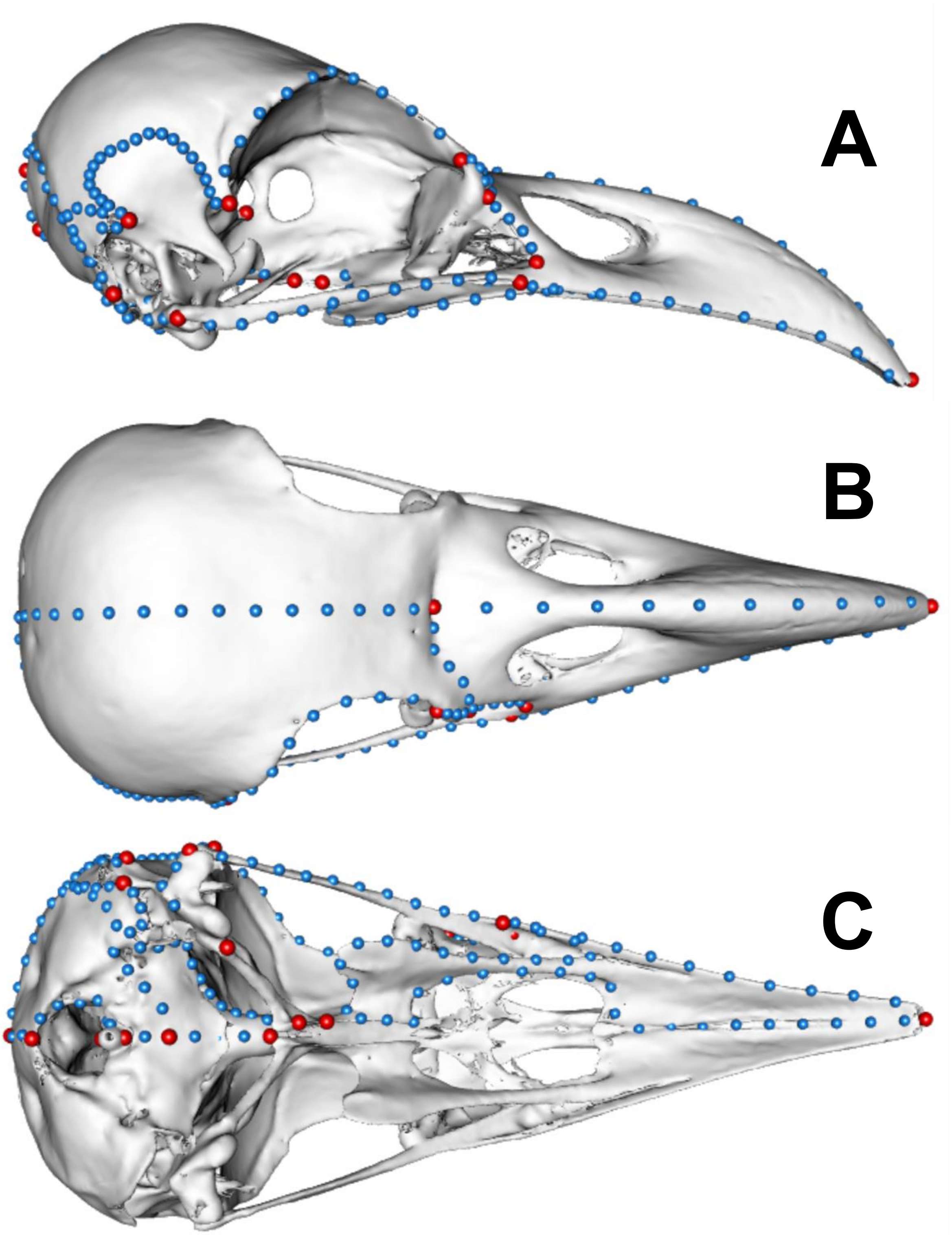


**Figure S2: Landmark layout on skull of *Corvus albus*.** Right lateral (A), dorsal (B) and ventral (C) views are shown. Red points represent anatomical (Bookstein type I and II) landmarks; blue points represent semilandmark curves.

**Endocast**

**Anatomical landmarks**

1. Anterior point of olfactory bulb on midline
2. Posterior contact of cerebral hemispheres on midline
3. Posterior lateral contact between cerebrum and optic lobe
4. Anterior lateral contact between cerebrum and optic lobe
5. Anterior contact between optic lobe and medulla
6. Posterio-dorsal contact between optic lobe and midbrain
7. Anterior point of cerebellum on midline
8. Anterior point of lateral cerebellar ridge
9. Posterior point of lateral cerebellar ridge
10. Posterior point of cerebellum on midline
11. Anterior point of medulla on midline
12. Contact point between medulla, midbrain and optic lobe
13. Posterior-most lateral contact point between medulla and midbrain at margin of foramen magnum
14. Posterior-most point of medulla on midline, at margin of foramen magnum

**Semilandmark curves**

1. Lmks 1 – 2, along midline between cerebral hemispheres
2. Lmks 2 – 3, along posterior boundary of cerebrum
3. Lmks 3 – 4, along boundary of cerebrum and optic lobe
4. Lmks 4 – 1, along anterior margin of cerebrum
5. Lmks 4 – 5, along anterior boundary of optic lobe
6. Lmks 5 – 6, along ventral/posterior boundary of optic lobe
7. Lmks 6 – 3, along dorsal/posterior boundary of optic lobe
8. Lmks 7 – 8, along anterior margin of cerebellum
9. Lmks 8 – 9, along lateral margin of cerebellar ridge
10. Lmks 9 – 10, along dorsal boundary of cerebellum
11. Lmks 10 – 7, along dorsal midline of cerebellum
12. Lmks 11 – 12, along anterior margin of medulla
13. Lmks 12 – 13, along lateral boundary of medulla
14. Lmks 13 – 14, along contact margin with foramen magnum
15. Lmks 14 – 11, along ventral midline of medulla

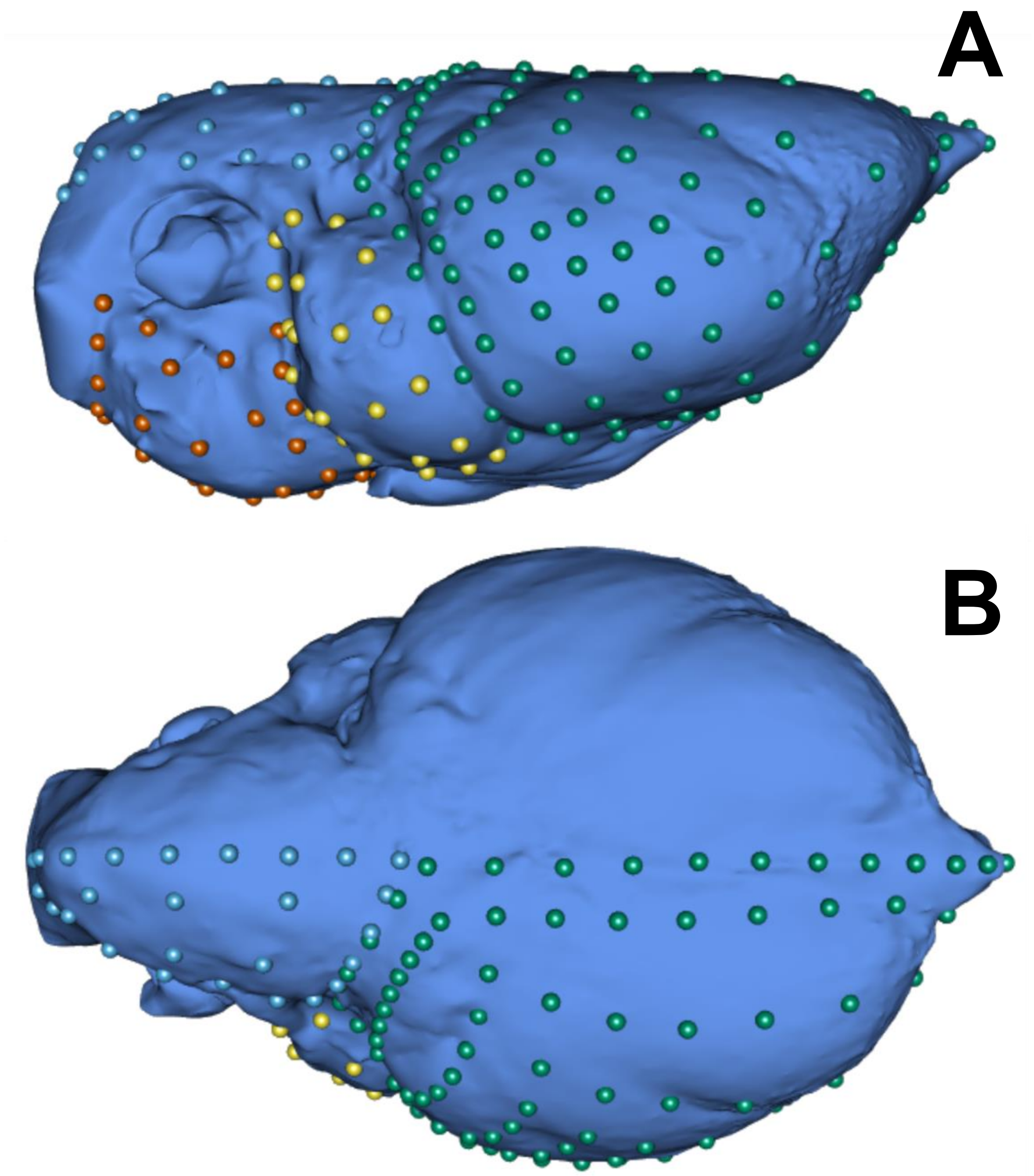


**Figure S3: Landmark placement of endocast, shown on endocast of *Anhinga anhinga.*** Right lateral (A) and dorsal (B) views are shown, with rostral end to the right. The four regions are shown in green (cerebrum), yellow (optic lobe), cerebellum (blue) and medulla (red).

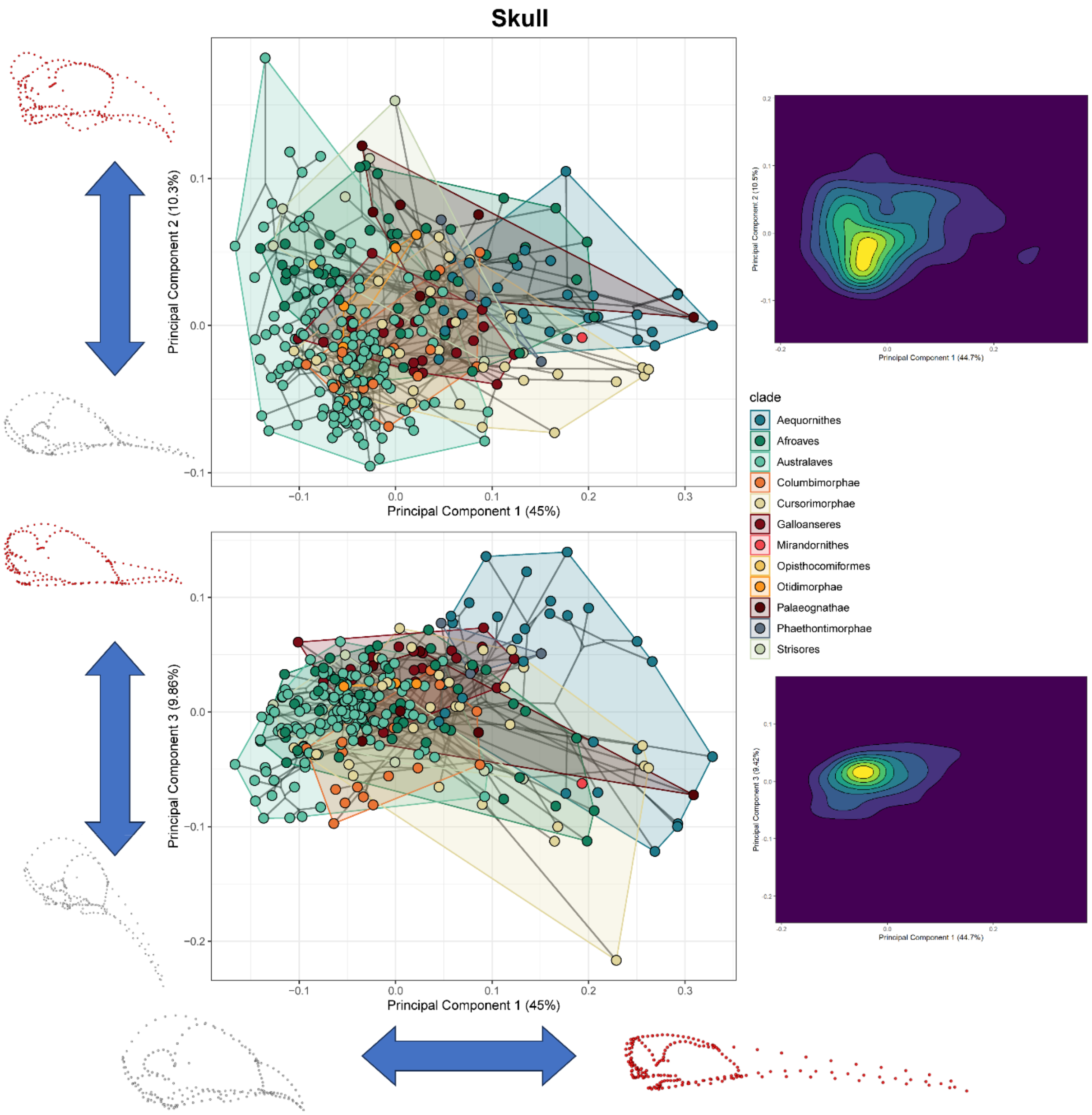


**Figure S4: Phylomorphospace of whole skull shape.** PCs 1 and 2 (upper plot) and PCs 1 and 3 (lower plot) are plotted with shape transformations shown adjacent to axes. Heat maps to left show density distributions of specimens of adjacent phylomorphospaces.

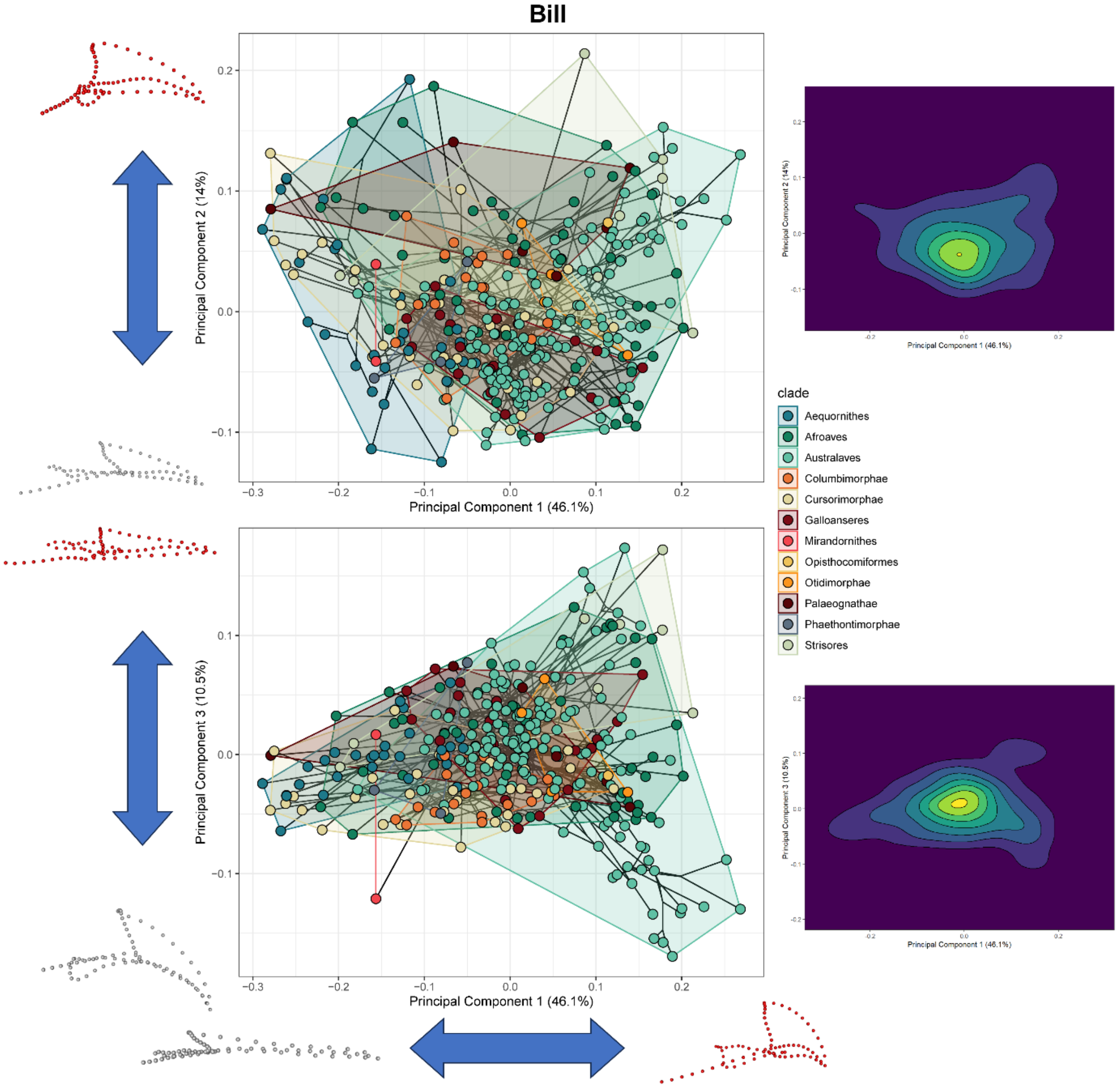


**Figure S5: Phylomorphospace of beak shape.** PCs 1 and 2 (upper plot) and PCs 1 and 3 (lower plot) are plotted with shape transformations shown adjacent to axes. Heat maps to left show density distributions of specimens of adjacent phylomorphospaces.

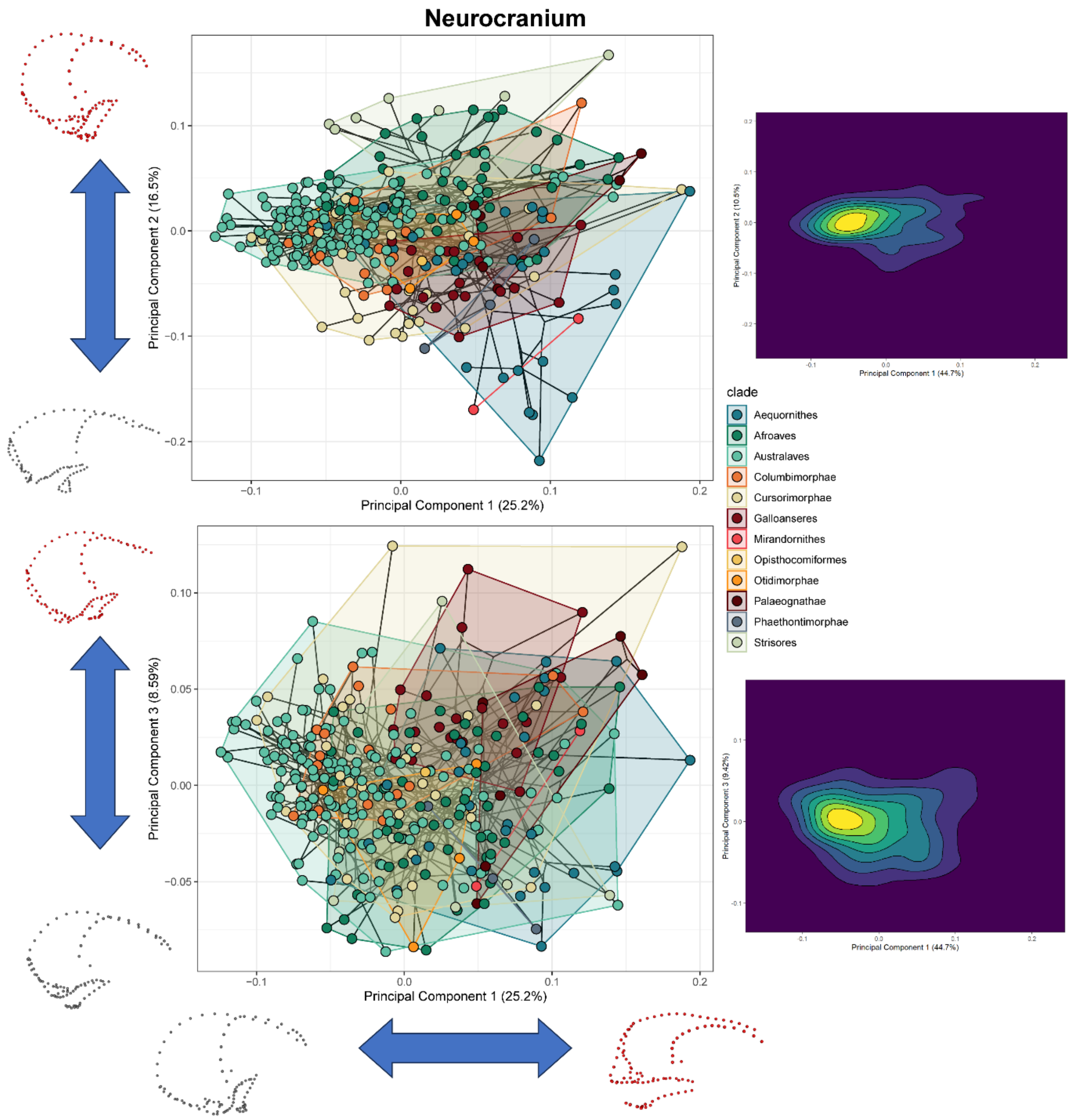


**Figure S6: Phylomorphospace of neurocranium shape.** PCs 1 and 2 (upper plot) and PCs 1 and 3 (lower plot) are plotted with shape transformations shown adjacent to axes. Heat maps to left show density distributions of specimens of adjacent phylomorphospaces.

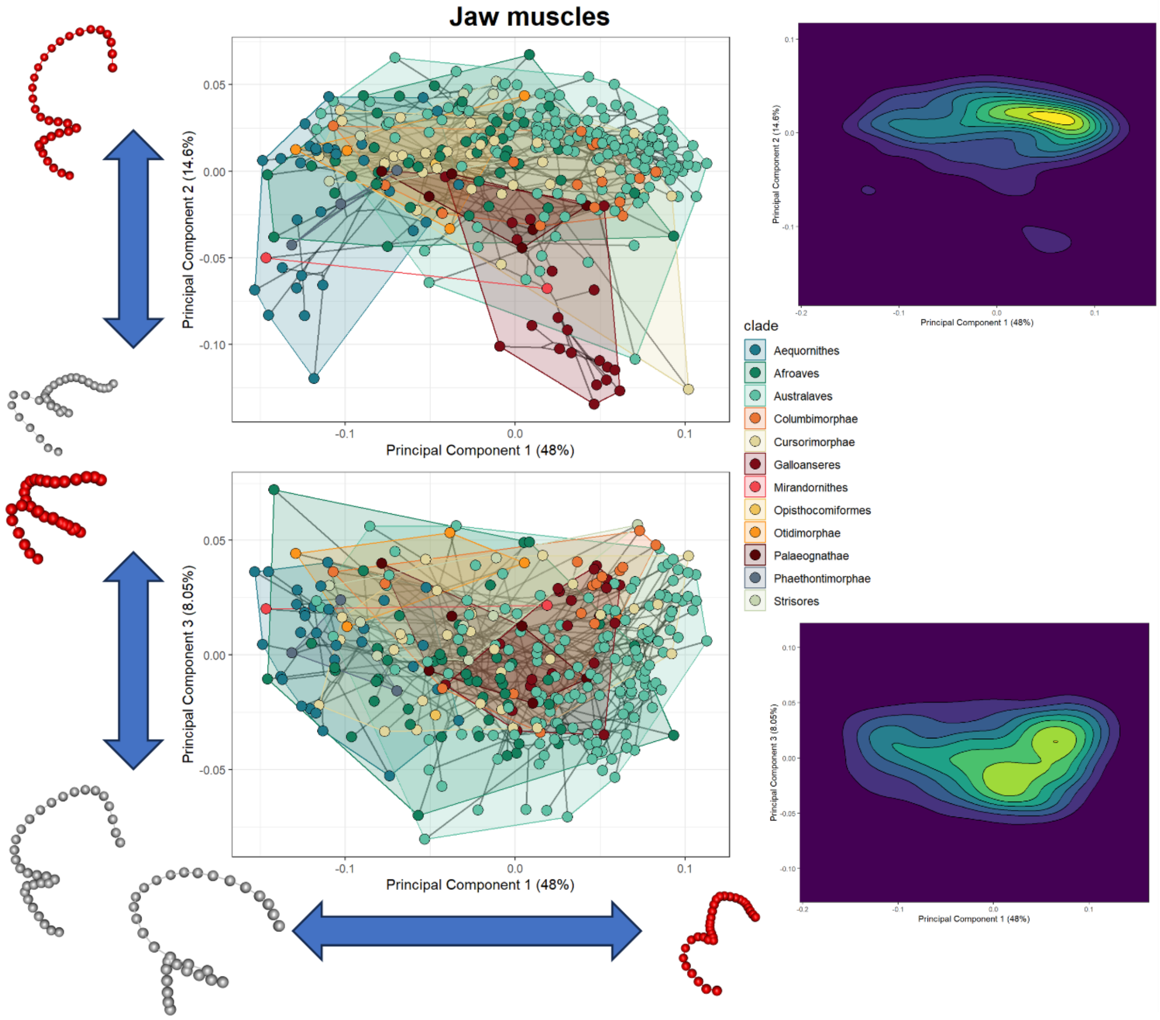


**Figure S7: Phylomorphospace of jaw muscle (adductor mandibulae and depressor mandibulae) shape.** PCs 1 and 2 (upper plot) and PCs 1 and 3 (lower plot) are plotted with shape transformations shown adjacent to axes. Heat maps to left show density distributions of specimens of adjacent phylomorphospaces.

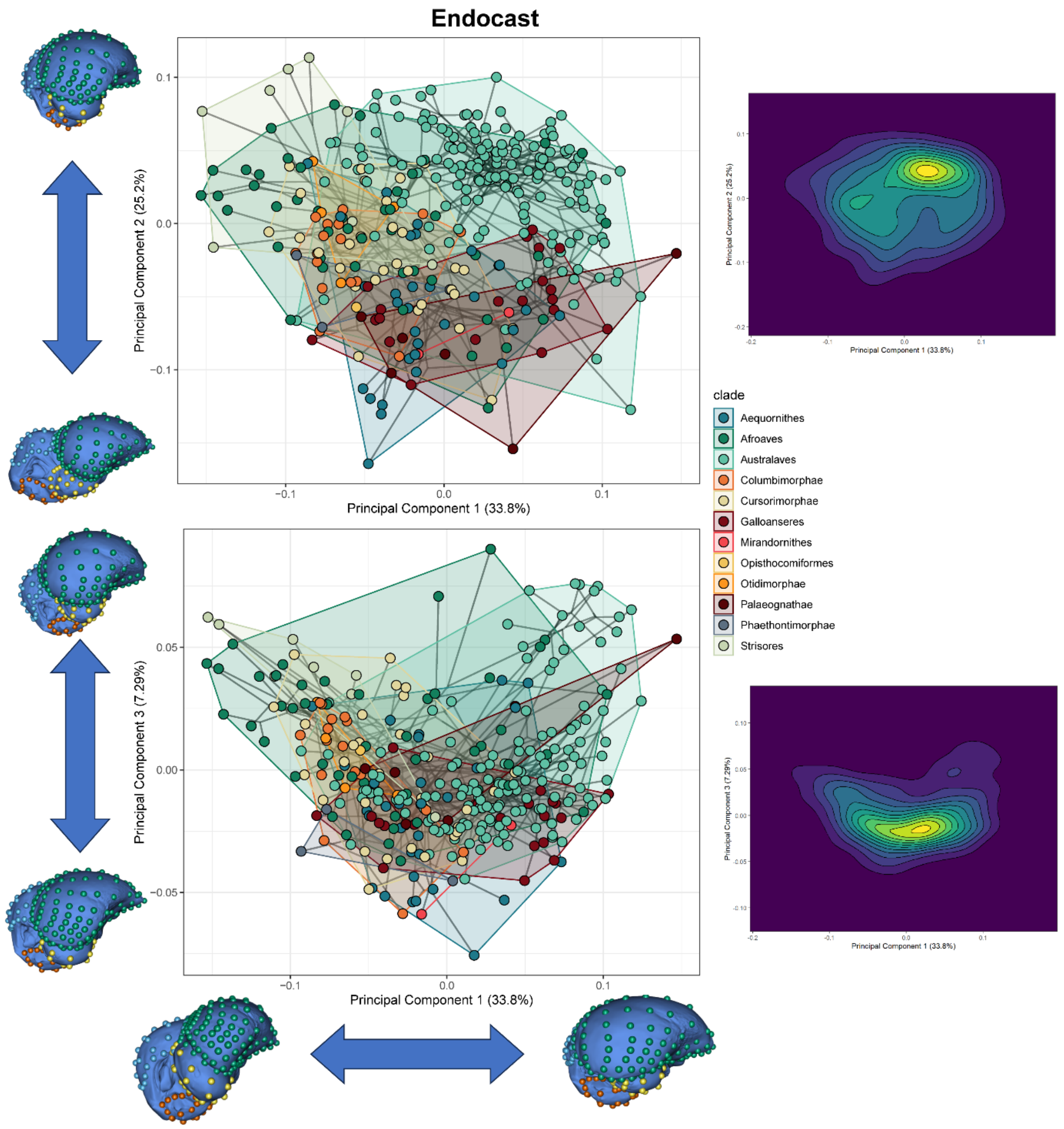


**Figure S8: Phylomorphospace of endocast shape.** PCs 1 and 2 (upper plot) and PCs 1 and 3 (lower plot) are plotted with shape transformations shown adjacent to axes. Heat maps to left show density distributions of specimens of adjacent phylomorphospaces.

**Table S1: Results of 2-block PLS analyses.** Pairwise results are shown for r-PLS (upper triangle) and effect size (lower triangle). All results are significant (p = <0.05).

| | Endocast | Beak | Neurocranium | Jaw muscles | Residual orbit diameter |
|---|---|---|---|---|---|
| **Endocast** | | 0.376 | 0.771 | 0.540 | 0.452 |
| **Beak** | 3.961 | | 0.582 | 0.421 | 0.341 |
| **Neurocranium** | 9.621 | 7.561 | | 0.667 | 0.642 |
| **Jaw muscles** | 6.038 | 4.480 | 9.204 | | 0.293 |
| **Residual orbit diameter** | 4.278 | 3.758 | 7.461 | 3.144 | |

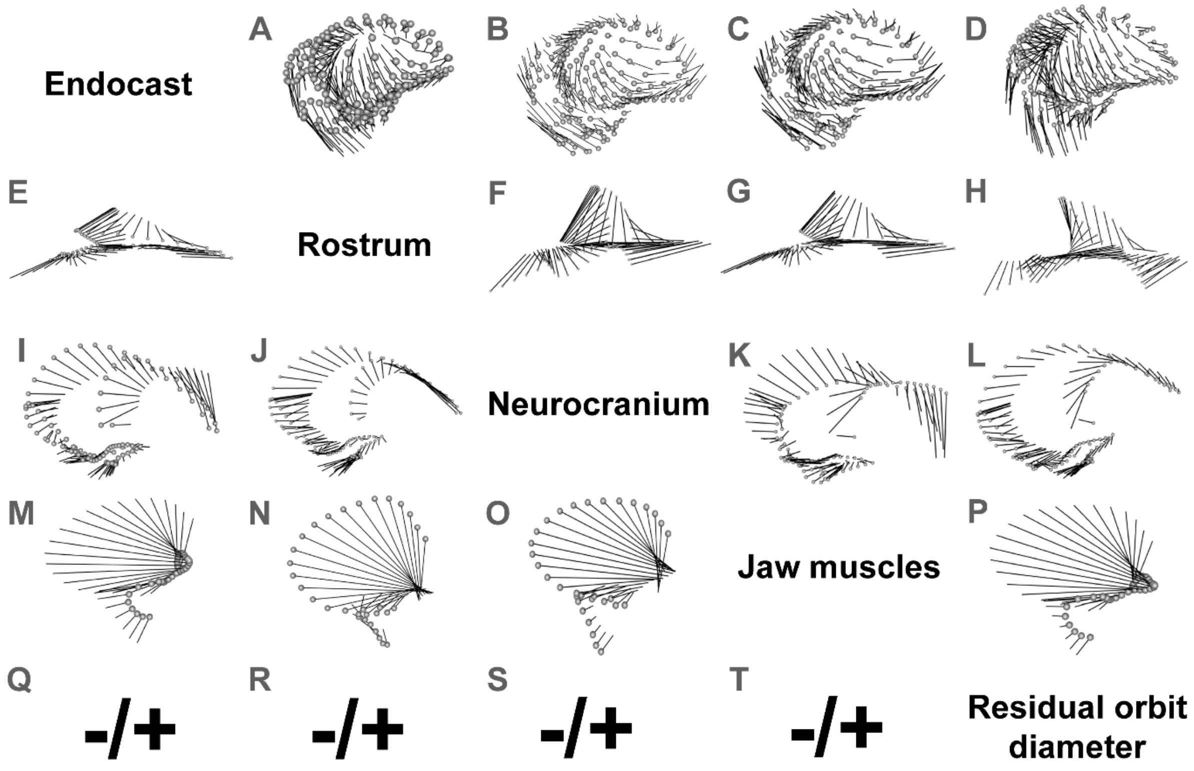


**Figure S9: Shape correlates of two-block PLS analyses.** Traits are read horizontally and pairwise interactions are read at the vertical intersection with the relevant trait (endocast A – D; Rostrum E – H; neurocranium I – L; jaw muscles M – P; residual orbit diameter Q – T). Shapes are shown at minimum (grey spheres) to maximum values, with black lines representing transformation vectors for each landmark.

**Table S2: Outline of seven models tested with mvSLOUCH** (Bartoszek et al., 2012 & 2024)**. A**-matrices correspond to models, with rows and columns corresponding to traits of interest. Interactions are inferred by columns influencing rows. Diagonals are assigned to "+", meaning that there is positive selection on that trait and that strength of selection is estimated by the model. Values that are assigned "0" means that the entry is constrained to be 0 in the estimation procedure (i.e. no interaction between traits). Interactions marked "?" indicates that values are free to vary in the estimation procedure, and implying interaction between two traits that is to be estimated. Traits are abbreviated to jaw muscle shape (MS), residual eye size (RES), endocast shape PC1 and PC2 (BS1 and BS2, respectively), neurocranium shape (NS), and beak shape (RS). Model numbers correspond to **1**: Hand in Glove 1; **2**: Hand in Glove 2; **3**: Spatial Packing 1; **4**: Spatial Packing 2; **5**: Spatial Packing 3; **6**: Functional Matrix 1; **7**: Functional Matrix 2; **8**: Ecological Selection 1; **9**: Ecological Selection 2; **10**: Ecological Selection 3; **11**: Modular; **12**: Fully Integrated. See Table 1 for graphical representations of interactions.

| Model | A | MS | RES | BS1 | BS2 | NS | RS |
|---|---|---|---|---|---|---|---|
| 1 | MS | + | 0 | 0 | 0 | ? | 0 |
| | RES | 0 | + | 0 | 0 | ? | 0 |
| | BS1 | 0 | 0 | + | ? | 0 | 0 |
| | BS2 | 0 | 0 | ? | + | 0 | 0 |
| | NS | 0 | 0 | ? | ? | + | 0 |
| | RS | 0 | 0 | 0 | 0 | ? | + |
| 2 | MS | + | 0 | 0 | 0 | ? | 0 |
| | RES | 0 | + | 0 | 0 | ? | 0 |
| | BS1 | 0 | 0 | + | ? | ? | 0 |
| | BS2 | 0 | 0 | ? | + | ? | 0 |
| | NS | ? | ? | ? | ? | + | ? |
| | RS | 0 | 0 | 0 | 0 | ? | + |
| 3 | MS | + | 0 | ? | ? | 0 | 0 |
| | RES | 0 | + | ? | ? | 0 | 0 |
| | BS1 | 0 | 0 | + | ? | 0 | 0 |
| | BS2 | 0 | 0 | ? | + | 0 | 0 |
| | NS | 0 | 0 | ? | ? | + | 0 |
| | RS | 0 | 0 | ? | ? | 0 | + |
| 4 | MS | + | 0 | ? | ? | 0 | 0 |
| | RES | 0 | + | ? | ? | 0 | 0 |
| | BS1 | ? | ? | + | ? | ? | ? |
| | BS2 | ? | ? | ? | + | ? | ? |
| | NS | 0 | 0 | ? | ? | + | 0 |
| | RS | 0 | 0 | ? | ? | 0 | + |
| 5 | MS | + | 0 | 0 | 0 | 0 | ? |
| | RES | 0 | + | ? | ? | 0 | 0 |
| | BS1 | 0 | 0 | + | ? | 0 | 0 |
| | BS2 | 0 | 0 | ? | + | 0 | 0 |
| | NS | 0 | 0 | ? | ? | + | 0 |
| | RS | 0 | 0 | ? | ? | 0 | + |
| 6 | MS | + | 0 | 0 | 0 | 0 | 0 |
| | RES | 0 | + | 0 | 0 | 0 | 0 |
| | BS1 | 0 | 0 | + | ? | 0 | 0 |
| | BS2 | 0 | 0 | ? | + | 0 | 0 |
| | NS | ? | ? | ? | ? | + | ? |
| | RS | 0 | 0 | 0 | 0 | 0 | + |
| 7 | MS | + | ? | ? | ? | 0 | ? |
| | RES | ? | + | ? | ? | 0 | ? |
| | BS1 | ? | ? | + | ? | 0 | ? |
| | BS2 | ? | ? | ? | + | 0 | ? |
| | NS | ? | ? | ? | ? | + | ? |
| | RS | ? | ? | ? | ? | 0 | + |
| 8 | MS | + | 0 | 0 | 0 | ? | 0 |
| | RES | 0 | + | 0 | 0 | ? | 0 |
| | BS1 | 0 | 0 | + | ? | 0 | ? |
| | BS2 | 0 | 0 | ? | + | 0 | ? |
| | NS | 0 | 0 | ? | ? | + | 0 |
| | RS | 0 | 0 | 0 | 0 | 0 | + |
| 9 | MS | + | 0 | 0 | 0 | ? | 0 |
| | RES | 0 | + | 0 | 0 | ? | 0 |
| | BS1 | 0 | 0 | + | ? | ? | 0 |
| | BS2 | 0 | 0 | ? | + | ? | 0 |
| | NS | 0 | 0 | 0 | 0 | + | ? |
| | RS | 0 | 0 | 0 | 0 | 0 | + |
| 10 | MS | + | 0 | 0 | 0 | 0 | ? |
| | RES | 0 | + | 0 | 0 | ? | 0 |
| | BS1 | 0 | 0 | + | ? | 0 | ? |
| | BS2 | 0 | 0 | ? | + | 0 | ? |
| | NS | 0 | 0 | ? | ? | + | 0 |
| | RS | 0 | 0 | 0 | 0 | 0 | + |
| 11 | MS | + | 0 | 0 | 0 | 0 | ? |
| | RES | 0 | + | ? | ? | ? | 0 |
| | BS1 | 0 | ? | + | ? | ? | 0 |
| | BS2 | 0 | ? | ? | + | ? | 0 |
| | NS | 0 | ? | ? | ? | + | 0 |
| | RS | ? | 0 | 0 | 0 | 0 | + |
| 12 | MS | + | ? | ? | ? | ? | ? |
| | RES | ? | + | ? | ? | ? | ? |
| | BS1 | ? | ? | + | ? | ? | ? |
| | BS2 | ? | ? | ? | + | ? | ? |
| | NS | ? | ? | ? | ? | + | ? |
| | RS | ? | ? | ? | ? | ? | + |
